# Age-related Macular Degeneration is associated with faster rates of structural brain changes and widespread differences in connectivity

**DOI:** 10.1101/2022.07.19.500546

**Authors:** Jacques A Stout, Rui Dai, Robert J Anderson, Scott Cousins, Jie Zhuang, Eleonora M Lad, Diane Whitaker, David Madden, Guy Potter, Heather E Whitson, Alexandra Badea

## Abstract

Age-related macular degeneration (AMD) is a prevalent disease impeding vision. More recently, AMD has also been linked to cognitive impairment, such as deficits in language and memory skills. In order to better understand the extent of AMD-related changes in the whole brain structure and connectivity, we have conducted an MRI diffusion acquisition study on 40 participants (20 diagnosed with AMD and 20 controls). These acquisitions were then performed again in a follow up two years later. We developed novel analysis methods for diffusion based tractography and connectomes to better determine which, if any, brain region connections saw the greatest changes between the AMD and the age-matched control groups. Using voxel-based analysis, we identified atrophy in AMD participants in the cuneate gyrus, which has been associated with vision, and the left superior temporal gyrus, which has been associated with language, while later acquisitions compounded this with a deficiency in the bilateral cingulate gyrus, itself linked to higher cognition and memory. These regional atrophy findings support that people with AMD experience widespread neuronal degradation that is not limited to retinal neurons. Regions that saw drastically lowered fractional anisotropy among AMD vs. control included the visual cortex, such as the bilateral occipital lobe and the frontoparietal cortex. Tensor Network Principal Component Analysis (TN-PCA) isolated lingual and temporal connections as important differentiators of AMD connectomes compared to controls, thus supporting our morphometric and texture findings in regions related to vision, but also connectopathies of language and memory brain regions. Bundle based analyses in baseline data revealed that the lingual gyrus had greater spread of tracts overall in the AMD participants, which may be explained by prior reorganization in this area, demonstrating a connection between retinal health and lingual structure. Moreover, we noted group differences in the interhemispheric temporal connections, and lingual cerebellar connections, supporting extensive downstream effects of vision loss. Our bundle-based analyses expand the toolset available for neuroimaging-based phenotyping, and reveal widespread changes in AMD participants beyond brain regions and tractography networks directly involved in vision processing, including those involved in language and memory.

## Introduction

Age-related macular degeneration (AMD) is one of the most common causes of legal blindness in older adults of developed countries [1, 2], affecting nearly 200 million individuals and up to 300 million by 2040 [3]. Previous studies have found that AMD is associated with greater cognitive decline compared to control populations without AMD [4]. However, the neural pathways behind the associated decline have not been well-delineated, and further understanding of the pathological structural changes may provide greater insights into the underlying mechanisms.

AMD is a debilitating disease of the eye with progressive deterioration of photoreceptors, retinal pigment epithelium, Bruch’s membrane, and the choroid [5]. Despite the availability of stabilizing treatments, damage to vision is progressive and irreversible, often resulting in legal blindness. Early and intermediate AMD are characterized by different quantities of small, medium, and large-sized drusen. Advanced or late AMD is separated into two categories: wet (vascular) or dry (nonvascular) AMD. Dry AMD is characterized by drusen and atrophy extension to the center of the macula. Wet AMD is distinguished by choroidal and retinal neovascularization, which results in subretinal fluid, lipid deposition, hemorrhage, retinal pigment epithelium detachment, and fibrotic scars [6].

Many findings suggest that AMD not only affects vision but is associated with changes in cognitive performance [4, 7–10]. Participants with AMD suffer greater cognitive decline, at a faster rate, with lower results on standardized tests based on Digit Symbols and Word Fluency than their peers without early AMD. Research also indicates that this decline is particularly strong for tasks of phonemic verbal fluency, independent from visual ability [7, 11, 12]. Using cross-sectional data from the same cohort as that used by this very study, it was shown that AMD patients, compared to peers with normal vision, exhibit specific brain changes involving language areas and connectivity[13]. Meanwhile, other studies have suggested that AMD-associated differences in brain structure may be partially compensated for by important functional restructuring [14, 15]. It is not known whether AMD causes changes to the brain (directly or indirectly), or whether specific brain changes may commonly co-occur with AMD, perhaps due to shared risk factors for other neurodegenerative conditions. This question cannot be answered definitively in the absence of prospective, randomized data (a design that is impossible for obvious ethical and practical reasons). Nonetheless, it is important to understand specific brain and cognitive changes associated with AMD, especially as new therapies emerge for dementia prevention and treatment in high risk populations[16–18].

Studies have demonstrated that there is a significant association between risk of AMD and Alzheimer’s disease compared to controls [19–21]. In AMD, degeneration of neuro-retina is associated with accumulation of drusen, which is primarily composed of extracellular beta-amyloid and lipids [22–25]. Participants with Alzheimer’s disease also have a higher incidence of drusen deposition in the retina compared to controls. Both AMD and Alzheimer’s disease demonstrate histopathological accumulation of beta-amyloid over time and are associated with microangiopathic vascular changes and local inflammation prior to clinical appearance of pathology [26–29].

One plausible mechanism by which AMD could contribute to structural brain changes, superimposed on normal age-related cognitive decline, is through deprivation of sensory input caused by loss of vision. In participants with dual hearing and vision loss, there is a significantly increased risk of both developing Alzheimer’s disease and non-specific cognitive decline and dementia compared to a control population with normal sensory function[30–32]. For most studies, this effect appears to be more important for those with visual than hearing impairment [31, 33] though some indicated the opposite[34].

MRI studies of participants with hearing loss found that hearing impairment was associated with smaller brain volumes in the primary auditory region, but not with other regions [35]. This suggests that typical cognitive decline due to sensory deprivation will be specific to the brain regions associated with the deprived sensations. In contrast, if AMD has an effect on cognitive decline, it is a strong indicator that its effects are not limited to the primary visual cortex but also affects other regions, though more evidence is needed to understand this connection. Previous studies have identified connections related to cognitive decline in this cohort but other factors, such as aging and AMD may affect connections between remote brain areas [13]. Here, we used voxel-based analyses for volume and texture based on diffusion tensor imaging, where fractional anisotropy characterizes microstructural integrity and allows to reconstruct brain connectivity networks based on tractography.

Multiple studies have demonstrated that participants with AMD showed atrophy of brain regions that were linked to vision such as the left optic tract [13] and the visual cortex [14]. However we have also observed that there is a stronger relation of cognitive performance with specific functional networks and connections among individuals with AMD [13, 36]. A primary objective of this study was to examine brain changes over a time span of two years, to determine whether AMD patients, compared to controls, exhibited a faster rate of volume loss or white matter decline, or whether the group differences observed at baseline merely persisted. A second objective was to identify specific pathways that change over time in the context of AMD and the extent of morphometric and microstructural changes in order to provide a better understanding of the mechanisms of both the visual decline as well as the cognitive changes associated with AMD. Furthermore, using novel tractography analyses, we explored the evolution of specific region connections of interest, and whether connectivity differences between AMD and control participants appear or increase after a longer period of time post-diagnosis. Finally, we conducted a novel connectome analysis in baseline data to determine which regions demonstrated the greatest AMD-related structural differences and their evolution over time. The detection of wide spread group differences affecting regions and connections involved in cognitive decline will help better understand the mechanisms of AMD-related cognitive deficits and can possibly be generalized to other neurodegenerative conditions.

## Methods

### Data Availability Statement

Datasets with limited personal health information can be made available via a request to the Authors. Receipt of any datasets will require a formal data sharing agreement between Duke University and the recipient, a brief protocol, and approval from the requesting researcher’s local Institutional Review Board. The code used in this analysis is available at ‘https://github.com/JacquesStout/DTC’ and any additional information needed can be provided through contact with the lead author.

### Participants

A total of 39 AMD (67% female, age range 61-94 years old (y.o.) mean age 75.7 y.o., standard deviation 8.8 years) and 33 control (61% female, age range 56-85 y.o., mean age 74.1 y.o., standard deviation 7.3 years) participants were recruited. Patients with AMD were referred from the Duke University Eye Center; age-matched controls were recruited from the friends and family of participants with AMD and from recruitment registries maintained in the Duke Aging Center. The inclusion criteria for AMD patients required participants to be over 50 years old and have a prior clinical diagnosis of either dry or wet AMD in one or both eyes causing visual impairment (20/40 or worse in the affected eye) for at least 1 year. All potential participants (AMD and controls) underwent exam by an ophthalmologist or optometrist to confirm AMD presence or absence and to exclude individuals with secondary ocular conditions (e.g., cataracts, glaucoma) causing uncorrectable vision impairment. Controls had lens-corrected vision of better than 20/40 in both eyes. Participants had to be able and willing to undergo MRI (no MRI-incompatible prosthesis, pacemaker, no current pregnancy, etc.), right-handed, and willing to return 2 years later. Individuals who lacked capacity to provide consent or had a documented diagnosis of dementia were excluded. Of the initial 72 participants recruited, 20 controls and 20 AMD participants returned 2 years later and were scanned for the second part of the experiment. The 40 individuals who provided 2-year MRIs were younger on average compared to the 32 who were lost to follow-up. This study exclusively used the data from the 40 individuals present in both the initial scan and at the follow up, to prevent any bias between the two scans. Most common reasons for lack of follow-up in this aging cohort were decline in general health status, death, new exclusions to MRI, or having moved from the area. All participants signed informed consents before the start of the study, per approval by the Duke University Medical Center Institutional Review Board.

### Imaging

Both anatomical and diffusion images were acquired from all participants. The anatomical images served to produce a brain parcellation through atlas based segmentation and were acquired using a 3D FSPGR [37] sequence whose parameters included: TR = 8.156ms; TE = 3.18ms; TI = 450ms; FOV = 25.6cm^2^, flip angle = 12°; voxel size = 1 x 1 x 1mm; 166 contiguous slices, sense factor = 2.

The diffusion scans served to provide morphometric and texture information, primarily for tractography and connectomics. They were acquired with 2D Spin-Echo / Echo-Planar imaging sequences. The parameters included: TR = 9,000ms, TE = 85.6ms; FOV = 25.6cm^2^; flip angle = 90°; voxel size = 1 x 1 x 2mm; 68 slices parallel to the AC-PC plane, 30 diffusion-weighted directions, b = 1000x/mm^2^, 4 non-diffusion-weighted reference images.

### Image Analysis

The diffusion images were preprocessed as described previously [38] here with additional denoising via Principal Component Analysis denoising [39]. Additionally, we used BET [40] for brain masking. The co-registration, eddy current correction, and the creation of diffusion parametric maps (fractional anisotropy) were created using software tools ANTs [41], DSI Studio [42], and DIPY [43].

All anatomical T1 subject images were registered to the IIT human brain atlas [44] using the SAMBA [45] pipeline applying a series of translation, rigid, and affine transformations. A minimum deformation template was generated, to reduce individual subject biases. All participants were mapped into the space of the minimum deformation template. The same registration was concurrently applied to the diffusion parameter maps previously obtained, and the reverse registration was used to brain the atlas labels into the original subject space, for region-based morphometry and texture analyses. Voxel-based analyses used the SPM toolbox [46] with a cluster-based False Discovery Rate (FDR) correction [47] and were used to compare volumetric and fractional anisotropy (FA) changes in the images.

The streamlines were reconstructed from the diffusion images using the Q-ball constant Solid Angle Reconstruction [48] applied via DIPY [43], with the tractography results built using a deterministic maximum direction getter. Other parameters included a relative peak threshold of 0.5, minimum separation angle at 25°, and binary stopping criterion using the brain mask. Streamlines were registered into the same space of the minimum deformation template using the same rigid and affine transforms as applied to the corresponding subject images.

The connectomes for the different diffusion images, defined as the matrix describing the number of white matter connections between different regions of the brain, were built using DIPY from the Q-ball model [48] tractography streamlines and using the IIT atlas, which defines 84 different gray matter regions [49].

There were several steps in our analysis. First the overall differences in the number of connections for all possible connection pairs between controls and AMD at a specific time point were compared directly to see whether there was a direct and easily observable increase or deficit of streamlines overall, for a given structure, and for a specific connection type. After the first broad test on 84×84 connection pairs, which, though specific, is a problem of high dimensionality. We subsequently ran a Tensor Network PCA (TN-PCA) [50] as described here [51], in order to reduce dimensions as this method has been shown to perform very well [52, 53] . TNPCA allows us to map the high-dimensional tensor network data to low-dimensional space. Together with grouping the initial and 2-year follow-up visit scans, this allowed us to determine differences between age-matched individuals with and without AMD, and longitudinal change (delta) in connectivity between these groups. Our results indicated which connections were most relevant for the different contrast comparisons.

The connections of interest (COI) determined by TNPCA to be the most discriminating between the controls and AMD participants were filtered using QuickBundles [54, 55] at a filter size of 3 mm, spatially matched after registering the fiber centroids with an affine transform and a warp, then sampled uniformly in 50 points. FA and Mean Diffusivity (MD) values were measured directly from the tractograms and FA images using DIPY. Overall fiber coherence to bundle was determined via FBC measurements [56] and bundle shape similarity was measured via two metrics. The first was purely distance based on the distance between bundles, and was determined by acquiring their centroids and calculating the minimum average direct-flip Euclidean distance between two bundles [55]. To complement this measurement, we also used the BUndle ANalytic values (BUAN) [57], which compare two bundles by also including the bundle adjacency and includes the shape similarity when computing the metric. The tracts and bundles statistical measurements were then all treated with R using two-tailed t-tests, with parametric values mapped to bundle centroids, and P-value, T-value evaluated. P-value < 0.05 was considered to be significant and the Cohen effect size was determined at the same time Cohen’s *d* = (*M*_2_ - *M*_1_) ⁄ *SD*_pooled_ where *SD*_pooled_ = √((*SD*_1_^2^ + *SD*_2_^2^) ⁄ 2) [58].

## Results

In this study, we have analyzed the brain morphometry (through the Jacobians of deformation fields) and texture (based on FA) using voxel-based analysis to reveal widespread morphometric and structural network connectivity changes in AMD participants, relative to controls over the span of two years.

### Volumetric Changes in AMD Participants

For volumetric comparison between AMD participants and controls, we analyzed the difference in brain volumes via Voxel Based Analysis (VBA) at the initial visit, two years later, and the rate of change between the two visits (**Figure 1**) with False Discovery Rate (FDR) correction. We found widespread volume decrease in multiple brain areas in AMD participants relative to matched controls. At the initial scan time, our results indicated lower volumes in participants with AMD especially in the left cuneate and lingual gyri [A], and bilateral superior and inferior temporal gyri [B]. The MRI scans at the second visit, two years after the first, indicate more extensive atrophy in the AMD group in the same regions as prior with new regions in the bilateral anterior cingulate [D] and superior frontal gyri. There were notable group differences, demonstrating that controls had a higher brain atrophy rates than the AMD participants in the right lateral occipital cortex and cuneate gyrus [C]. Comparison between the first and second visits revealed that the greatest group difference in rate of atrophy occurred in the cingulate gyrus [E] and the left temporal lobe [F].

**Figure 1:**
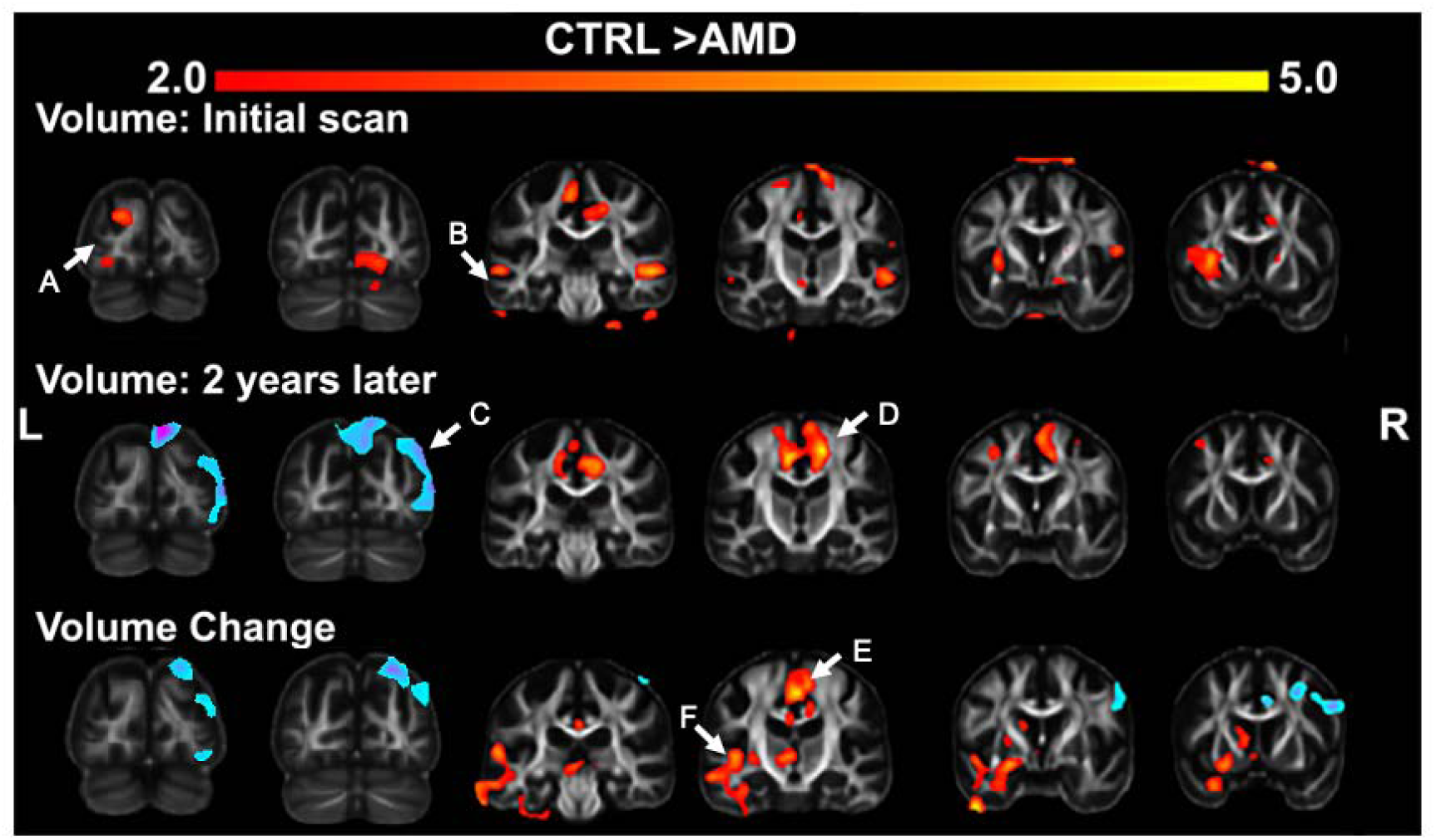
Most significant volumetric differences in Control group (CTRL) vs AMD group where CTRL > AMD as determined by VBA. First row: differences between groups at initial scan (FDR = 0.05) Second row: differences between groups two years later (FDR = 0.1). Third row: differences between rates of change in AMD versus controls (FDR = 0.2). Statistical maps represent FDR corrected anatomical maps at the most stringent thresholds, which were allowed to vary to yield comparable representations across the three rows), overlaid on the minimum deformation template. Red indicates CTRL > AMD, blue indicates CTRL < AMD.A) left cuneate (bottom) and left lingual gyrus (top) B) Inferior temporal gyrus (bottom left) and bilateral superior temporal gyrus (lower left and right) C) right lateral occipital cortex (right) and cuneate gyrus (top) D) bilateral anterior cingulate E) Cingulate gyrus F) Left Temporal lobe

### FA Changes in AMD Participants

For microstructural integrity analysis, we determined the most significant FA differences in the Control vs AMD patient groups during both acquisitions, as determined by VBA. Compared to the volumetric data, our results demonstrate a more selective pattern of group differences, with FA decline in AMD participants in the visual occipital cortices, as well as the anterior cingulate cortex, and frontoparietal cortex.

Our results at the initial time point indicate early texture impairment in inferior temporal lobe of participants with AMD compared to that of Control. For both the initial visit and the rates of change, the group differences did not survive FDR correction (uncorrected statistical maps are shown in **Supplementary Materials:** **Figure 1**). At the second visit however, we observed more extensive microstructural changes with additional involvement of the bilateral occipital lobe [A] and left frontoparietal cortex [C]. The rates of change indicated atrophy primarily in the left temporal lobe and frontal cortex which also did not survive FDR correction.

### Connectivity Changes in AMD Participants

After calculating the tractograms for each subject, the connectomes were estimated using atlas labels determined by SAMBA and grouped depending on whether the participants had AMD or were Control, and whether the scan was the initial scan or performed 2 years later.

Before examining a more specific comparison, AMD participants had on average slightly fewer connections than their Control counterparts overall (initial: CTRL 1.918×10^7^, AMD 1.875×10^7^; 2 year: CTRL 1.83×10^7^, AMD 1.690×10^7^). However, this difference was not significant (p-values: initial 0.74; 2 year 0.24), and disappeared entirely when comparing the initial AMD scans to the later Control scans. When looking at the initial scans, the connections with the biggest bias towards the Control group were local connections of the thalamic and sub-thalamic regions. While these differences were significant, given the high dimensionality of the connections, they did not survive an FDR correction. Meanwhile, comparison of the scans 2 years later showed a difference in the number of connections from the pericalcarine cortex to other regions (**Table 1**).

**Table 1.** Connections with highest T-Test results when directly comparing the number of connections in Control vs AMD participants, both during the initial visit and 2 years later. Result for AMD participants versus age matched controls for the initial visit and the second visit two years later.

|  | Connection | Number<br>streamlines<br>Control | Number<br>streamlines<br>AMD | T-values | p-values | FDR<br>corrected<br>p-values |
| --- | --- | --- | --- | --- | --- | --- |
| 1<sup>st</sup><br>Year | entorhinal R - Thalamus-Proper R | 3707 | 2275 | 3.19 | 1.30e-3 | 0.10 |
|  | entorhinal R – Hippocampus R | 28.499 | 31684 | 3.02 | 2.28e-3 | 0.17 |
|  | entorhinal R – Caudate L | 647 | 293 | 3.01 | 2.34e-3 | 0.17 |
|  | Parahippocampal R – Pallidum L | 355 | 168 | 2.81 | 4.40e-3 | 0.31 |
| After 2<br>years | pericalcarine L – lateral orbitofrontal L | 1263 | 595 | 3.83 | 1.81e-4 | 9.52e-3 |
|  | pericalcarine R – Pallidum L | 155 | 69 | 3.80 | 1.93e-4 | 0.013 |
|  | pericalcarine L – Pallidum L | 218 | 91 | 3.68 | 2.84e-4 | 0.019 |
|  | pericalcarine L -Pallidum R | 182 | 79 | 3.46 | 5.69e-4 | 0.038 |

To reduce the high dimensionality of our comparison, we next examined brain connectivity patterns with TN-PCA at the first visit, 2 years later, and the difference between the two time points (**Table 2**). Our results at the initial time point indicated that the region connections which had the greatest overall weight given the principal components used in differentiating the Control from the AMD participants were the lingual right to lingual left connection, followed by the fusiform right to superior temporal left, the superior frontal right to superior frontal left, and the inferior temporal right to superior temporal left (**Table 2A**).

**Table 2.**
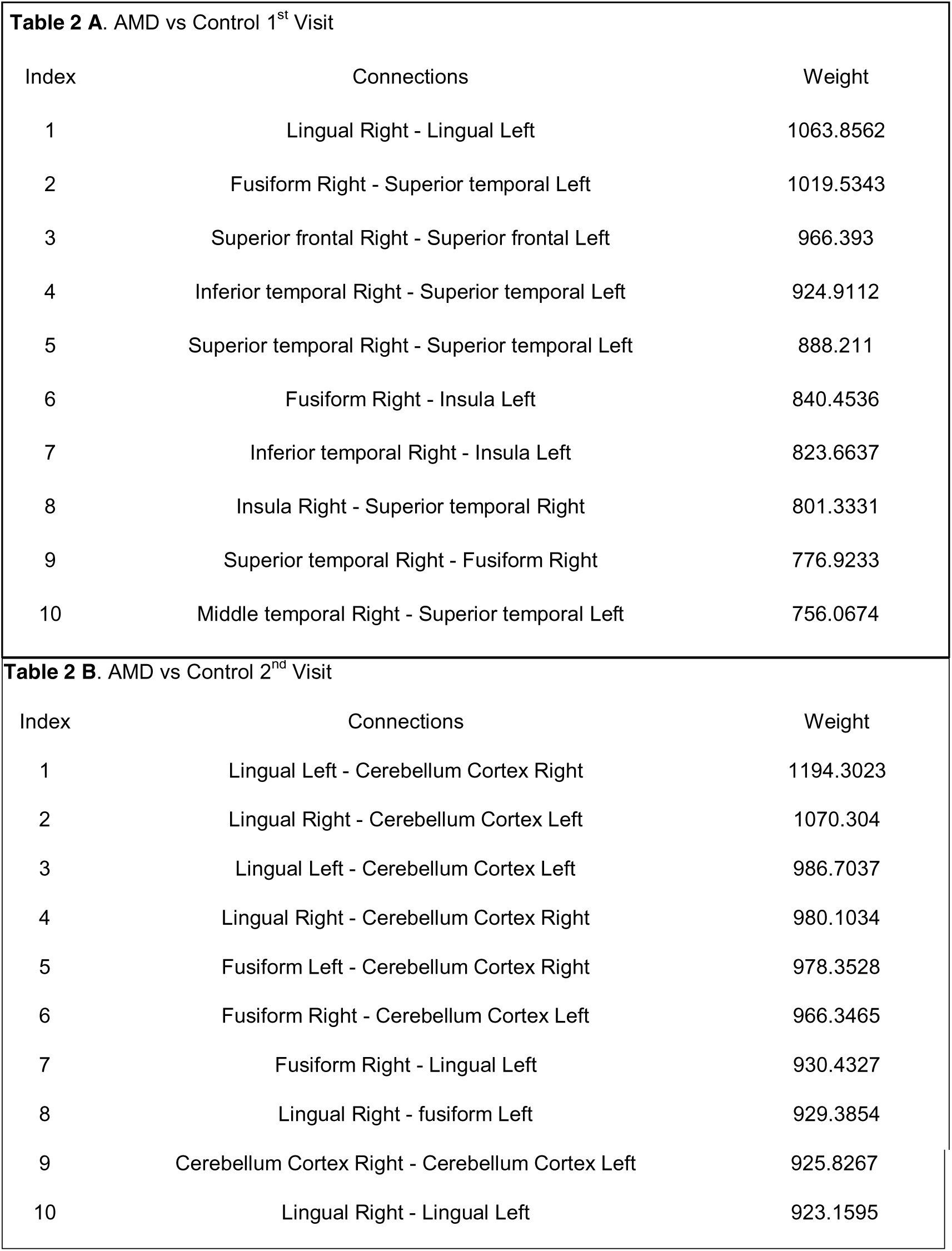
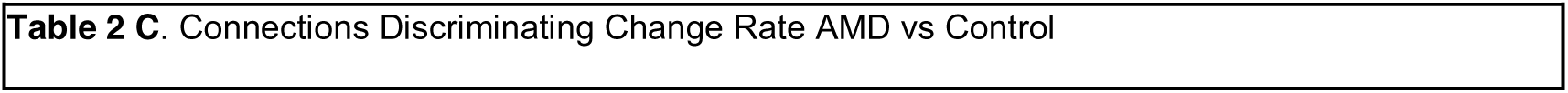
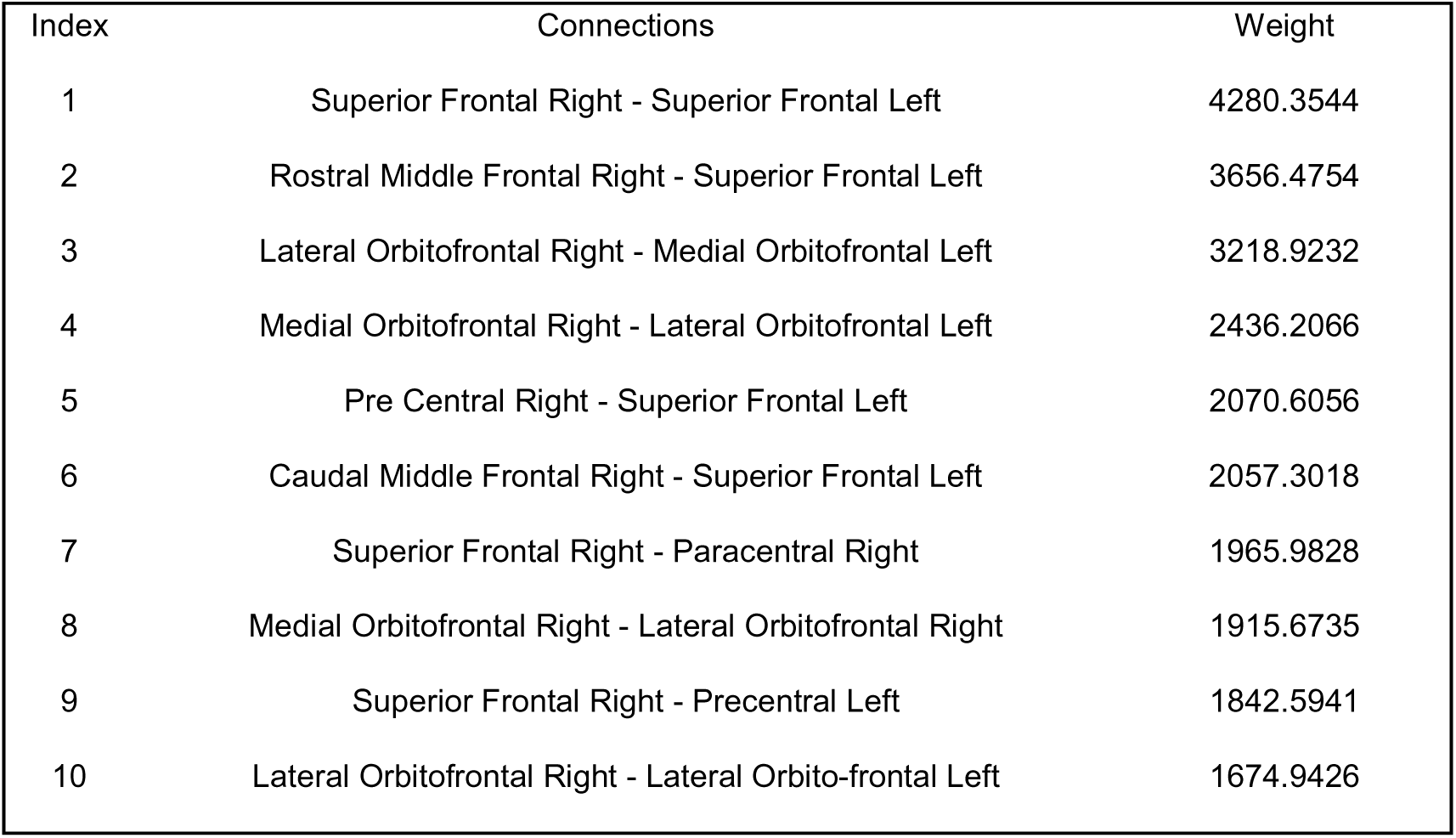
Top 10 TNPCA results for AMD participants versus age matched controls for: a) the initial visit; b) the second visit, two year later c) rate of change in connectivity

The same analysis was run with data obtained from the same participants 2 years later. This time, the connections that influenced our TN-PCA analysis most heavily were those corresponding to both the left and right side of the lingual gyrus to different cerebellum connections, with those going from the left to right having a heavier weight. Additionally, most of the following connection types involved either a part of the lingual region or the cerebellum (**Table 2B**).

When looking at the difference in the connectomes determined from participants from their initial visit to the one they had 2 years later, we found that the superior frontal left to the superior frontal right and to the rostral middle frontal right exhibited the largest group difference, with AMD demonstrating a greater loss of connections for these regions. Many of the other connectomes with significant group differences (AMD with greater loss than controls) involved the frontal regions or the lateral orbitofrontal and medial orbitofrontal regions (**Table 2C**).

Specific connections were selected from those that showed very high relevance during TN-PCA comparisons and unique regions forming part of the connection. These were: Lingual Right - Lingual Left (LinR-LinL), Inferior Temporal Right – Superior Temporal Left (InTempR-SuTempL), Lingual Left - Cerebellum Cortex Right (LinL-CerbR) and Lingual Right - Cerebellum Cortex Left (LinR-CerbL).

**Table 3** shows the direct comparisons of all the streamlines pertaining to each specific connection of Interest (COI) that originated from Control and AMD individuals, both on the initial analysis and 2 years later. We can see that due to the high number of streamlines, the p-values bring little information. However, the Cohen effect provides a more reliable narrative.

**Table 3.** Tractography data breakdown (Length of streamlines, FA found along streamlines) of all tracts associated to each connection type found in all AMD participants and in all Control participants.

|  | Connections | Group | Mean Length | p-val<br>T-val<br>Cohen | Mean FA | p-val<br>T-val<br>Cohen | p-val<br>T-val<br>Cohen |
| --- | --- | --- | --- | --- | --- | --- | --- |
| 1<sup>st</sup><br>Year | Lingual Right –<br>Lingual Left | Control | [42.6 - 44.1] | <0.01<br>-6.63<br>-0.11 | [0.117 -<br>0.119] | <0.01<br>-14.05<br>-0.23 | <0.01<br>3.25<br>0.05 |
|  |  | AMD | [46.4 - 48.2] |  | [0.130 -<br>0.133] |  |  |
|  | Inferior Temporal Right -<br>Superior Temporal Left | Control | [202.6 -<br>214.4] | <0.01<br>4.89<br>0.40 | [0.226 -<br>0.243] | <0.01<br>4.99<br>0.38 | 0.98<br>0.41<br>0.03 |
|  |  | AMD | [183.1 -<br>195.8] |  | [0.204 -<br>0.218] |  |  |
|  | Lingual Left –<br>Right Cerebellum Cortex | Control | [79.8 - 81.4] | <0.01<br>8.29<br>0.09 | [0.148 -<br>0.151] | 0.07<br>-1.80<br>-0.02 | <0.01<br>17.06<br>0.19 |
|  |  | AMD | [75.0 - 76.8] |  | [0.150 -<br>0.152] |  |  |
|  | Lingual Right –<br>Left Cerebellum Cortex | Control | [75.0 - 76.5] | <0.01<br>-7.94<br>-0.09 | [0.141 -<br>0.143] | <0.01<br>-7.81<br>-0.09 | <0.01<br>9.27<br>0.11 |
|  |  | AMD | [79.6 - 81.5] |  | [0.147 -<br>0.149] |  |  |
| After 2<br>years | Lingual Right –<br>Lingual Left | Control | [44.0 - 45.5] | 0.60<br>-0.52<br>-0.01 | [0.119 -<br>0.122] | <0.01<br>-21.74<br>-0.36 | <0.01<br>14.71<br>0.24 |
|  |  | AMD | [44.2 - 45.8] |  | [0.142 -<br>0.145] |  |  |
|  | Inferior Temporal Right -<br>Superior Temporal Left | Control | [199.6 -<br>209.8] | <0.01<br>4.16<br>0.34 | [0.248 -<br>0.264] | <0.01<br>9.75<br>0.79 | 0.29<br>1.07<br>0.09 |
|  |  | AMD | [184.1 - 195] |  | [0.200 -<br>0.213] |  |  |
|  | Lingual Left –<br>Right Cerebellum Cortex | Control | [74.7 - 76.1] | 0.06<br>1.85<br>0.05 | [0.150 -<br>0.152] | 0.57<br>0.56<br>0.01 | <0.01<br>5.50<br>0.01 |
|  |  | AMD | [73.7 - 75.1] |  | [0.145 -<br>0.152] |  |  |
|  | Lingual Right –<br>Left Cerebellum Cortex | Control | [77.9 - 79.4] | <0.01<br>8.48<br>0.10 | [0.153 -<br>0.143] | <0.01<br>13.00<br>0.15 | 0.22<br>-1.22<br>-0.01 |
|  |  | AMD | [73.1 - 74.6] |  | [0.143 -<br>0.145] |  |  |

The LinR-LinL streamlines showed higher FA among the AMD participants during the Initial Scan. This FA difference increased two years later, and the MD was also found to be higher in controls than AMD 2 years later even though this effect was fairly insignificant at the initial scan.

The streamlines linking InTempR-SuTempL at the initial scan showed a much higher Length and larger FA in Control groups than in those with AMD (Cohen Effect size: 0.40 / 0.38) even as the MD stayed almost the same. This difference in both Length and FA was still present two years later in the comparison of Control and AMD.

The differences in the Lingual – Cerebellum connections had a smaller Cohen effect size, but still showed some differences. Control group streamlines for LinL-CerbR had a greater length and MD than those found in AMD. However, the streamlines for LinR-CerbL for those same acquisitions found a greater length in AMD than Control.

When looking at the acquisitions taken 2 years later, the two interhemispheric connections (LinL-CerbR and LinR -CerbL) both had higher length and FA along the streamlines in Control than in the AMD acquisitions. The MD differences were relatively negligible.

**Figure 3** and **Figure 4** show the largest bundles created from the average tracts from both the Control group and the AMD group for the COI determined by the initial year TN-PCA and those found during the acquisitions taken 2 years later. The bundles resulting from the QuickBundles analyses within each group contain all those streamlines with similar shape and direction, and that are most frequently found in that specific group.

In the InTempR-SuTempL and to a lesser extent for LinR-LinL, we observe a higher spread in the observed bundles. There is a similar increase in the spread of the largest bundles across the template brain space to be found in LinR-CerbL, though it appears less pronounced. To better categorize these differences, we use Centroid distance and BUAN values to compare the different bundles which can all be seen in **Table 4**.

**Table 4.**
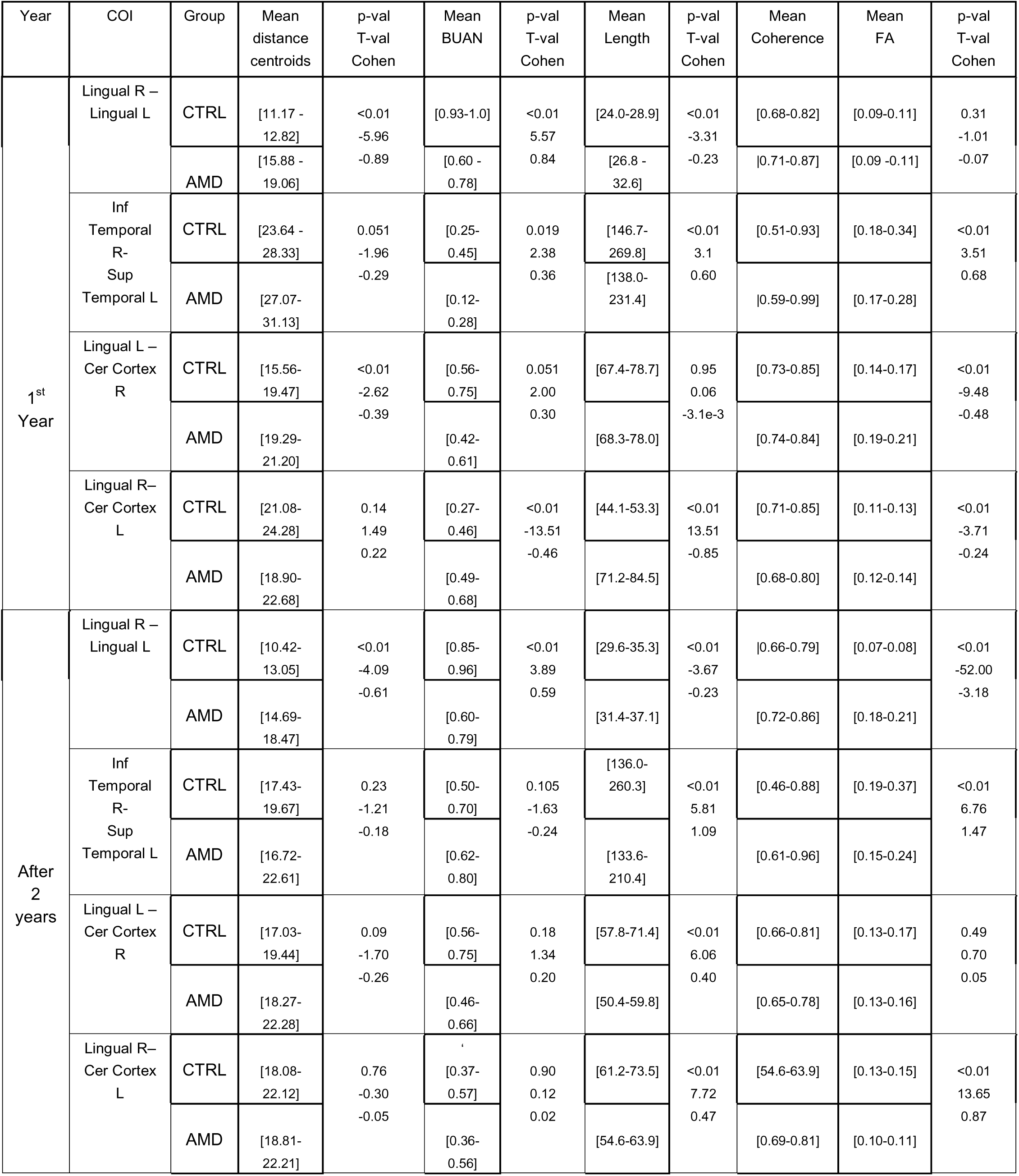
Similarity statistics for largest bundles and streamline statistics for streamline length, FA along the tracts

In **Figure 3**, the average FA along the streamlines found in the top bundles can be seen. The color mapping indicates that the streamlines associated to bundles seemed to have a higher FA in the Control group for the initial scan in the case of InTempR-SuTempL, as well as in LinL-CerbR and LinR-CerbL two years later.

**Figure 4** shows the same bundles of streamlines as **Figure 3**, however they are instead colored by the average coherence of the specific streamlines to the bundle that they belong to, i.e. their similarity to the centroids of the bundle itself. Overall, the LinR-LinL seems to have higher coherence in controls, however the group difference is less pronounced in InTempR-SuTempL. Other sets do not have enough of a difference between groups to be easily distinguishable visually.

While **Table 3** focused on the entirety of the streamlines from each COI, **Table 4** displays the statistics based on the ten largest bundles created from those same streamlines. This allows us to take a more in-depth look at the similarities of the streamlines of a given group when compared directly with each other. Of the top ten bundles created with a threshold of 3 mm in the overall sum of the streamlines found in 4 groups (Initial Control vs Initial AMD and 2year Control vs 2year AMD), we compared the centroids of each bundle and looked at the average distance of centroids from each other. We also calculated the BUAN score of the bundles. Both these metrics gives us a rough idea as to the similarity and proximity of the bundles to each other: a higher BUAN score and lower average distance for a specific group indicates that the bundles of a group corresponding to this connection tend to aggregate together.

Among the different COI, **Table 4** shows that the LinR-LinL connection has the closest aggregation of bundles overall. In controls, the LinR-LinL exhibited around 11-12 mm distance of centroids on average and around 0.9 BUAN; meanwhile, AMD bundles show a much greater level of dissimilarity, with 16-17mm distance and around 0.7 BUAN similarity. For both the Control and AMD groups, these values do not change in any substantial or significant way two years later. The FA of the LinR-LinL streamlines tended to be lower in controls than that found in AMD, with the latter being particularly noticeable for the 2 year later acquisition (0.07 => 0.20). Overall, the bundles for the Inferior Temporal Right to the Superior Temporal Left were closer together and had higher FA among the controls than those found in AMD individuals. Other results obtained for the Lingual regions to the Cerebellum did not show any notable trend.

**Figure 5** shows the largest bundles found in the Control group compared directly to those that are closest to them in space and shape, but found in AMD. The interhemispheric lingual connections showed a less clear trend, with two of the bundles showing higher FAs in controls, one showed similar patterns, while one bundle showed the reverse trend (**Figure 5A**). When directly comparing the largest bundles of the initial COI InTempR-SuTempL there was a clear tendency for higher FA among the Control group (**Figure 5B**).

At the second visit we noted for lower FA values among two of the bundles in the interhemispheric connections between the interhemispheric lingual -cerebellar connections, while one bundle showed the opposite trend. These results are suggestive of possible reorganization of the lingual-cerebellar connections.

## Discussion

AMD is one of the most significant causes of legal blindness in the developed world. Recent studies have found that AMD not only affects vision but is also associated with more severe decline in mental cognition compared to the general population [4, 7–10, 59]. The mechanisms behind the correlation are not well understood. In this study, we studied brain-wide morphometric, microstructural, and connectivity changes based on longitudinal diffusion weighted imaging (DWI) and identified widespread differences in brain networks among older adults with and without AMD, some of which may be linked to the faster cognitive decline compared to normal aging.

We obtained MRI scans from Control and AMD participants at two timepoints, approximately two years apart. The images were analyzed for differences in volume, FA, and rates of change, and we found important group differences in brain visual processing regions. Some of the differences increased over time. Microstructural changes as determined by fractional anisotropy were only noticeable for the second time point, 2 years after diagnosis.

During the initial scan, the cuneate and lingual gyri, as well as the superior and inferior temporal gyri all had greater volume in controls compared to participants with AMD. The cuneate and lingual gyri are known from other studies to be responsible for processing direct primary visual information of the occipital cortex [60–63]. Changes in these two gyri of the occipital cortex are expected in a patient with visual defect, as previous studies have demonstrated that participants with both early and late-onset blindness exhibit decreased occipital cortex volume [64].

Among participants who returned for a second visit two years later, the group difference in lingual gyri was no longer statistically significant. Instead, there was less volume atrophy of the lateral occipital cortex and cuneate gyrus in the AMD participants when compared to their Control counterparts, and after compensating for overall brain changes. This regional volumetric atrophy appeared in tandem with a nearby important decrease in FA in the posterior occipital lobe. This counter-intuitive result, particularly as it was not detected early, could potentially reflect the brain’s neuroplasticity, as loss of vision does not lead to permanent inactivation of the visual cortex, and there is a long-term reorganization with potential increase of brain activity in the occipital visual cortex long after an impairment in vision [65, 66].

However, both when looking at the acquisition taken 2 years later and when directly comparing the longitudinal changes of participants, the cingulate gyrus was shown to have important volumetric atrophy that was not significant during our initial analysis. The cingulate gyrus, a vital and complex part of the limbic system involved in the regulation of emotions and attention, exhibited important volumetric atrophy in participants with AMD. Sub-regions of the cingulate gyrus have all been associated with different types of higher brain function in previous studies. Damage to the anterior cingulate gyrus has been associated with apathy, akinesia, and increased stupor [67], while the posterior cingulate gyrus is involved in cognition, especially attention and internally directed cognition [68] and finally the superior frontal gyrus is associated with higher cognitive functions and working memory [69]. Due to these factors, it is possible that any observed cognitive deficiency observed in AMD could in fact be linked to the cingulate gyrus.

The left temporal lobe also saw fairly important longitudinal atrophy that was not symmetrical. This is of particular interest as this region includes Wernicke’s area, in the posterior third of the superior temporal lobe of the left hemisphere [70]. Language requires the coordination of several regions in the brain, most notably Wernicke’s area for speech interpretation and language conduction, and Broca’s area for speech production [71]. The corresponding location on the right hemisphere has also been noted to be important for word repetition tasks [72]. Given that AMD has been associated with poor performance on language-related cognitive tasks [12], it stands to reason that the brain changes reported here, including a rapid deterioration of volume and FA in the temporal lobes in patients with AMD compared to paired controls, could play a role in provoking accelerated cognitive decline.

The left frontoparietal cortex, which is associated with early cognitive decline [73] also exhibited markedly lower FA values in participants with AMD compared to controls at the same time point.

The superior and inferior temporal gyri are associated with integration of multimodal sensory processes. The inferior temporal cortex is known to be responsible for processing visual information from the occipital cortex [74]. The superior temporal cortex also has functions in visual and speech processing, especially for spatial-based and object-centered spatial-orientation reference systems [14, 75]. We hypothesize that the loss of structural integrity in this region may be reflected clinically as deterioration of language or semantic tasks over time in people affected by AMD.

The Voxel Based Analysis of FA seen in Figure 2 had overall fewer clear distinctions between Control and AMD that survived FDR correction. Only the inferior temporal lobe showed higher FA in controls. This region is important for verbal fluency, and impairment is associated with cognitive deficits that are seen in early Alzheimer’s disease [76]. The occipital lobes outlined in the second acquisition are the primary visual cortex, and changes in vision drives moderate changes to the cortex [77]. Meanwhile, the frontoparietal cortex that saw higher FA in controls is associated with higher-function conceptual action information and deficits result in spatial neglect [78–80].

**Figure 2:**
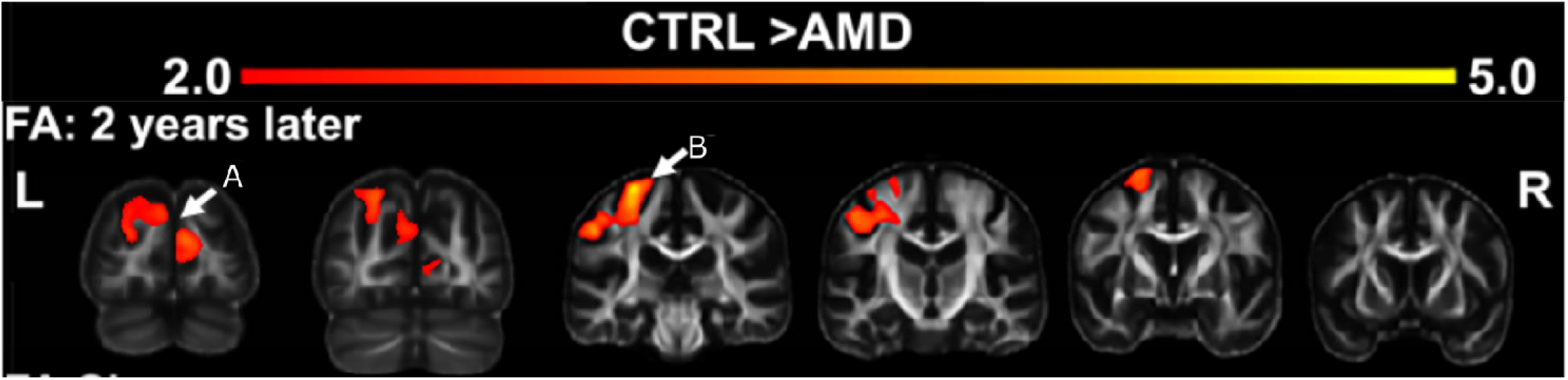
FA differences in Control group vs AMD group at the second visit (2 years after the initial visit) where CTRL > AMD as determined by VBA (FDR = 0.05). A) bilateral occipital lobe B) Left frontoparietal cortex

**Figure 3.**
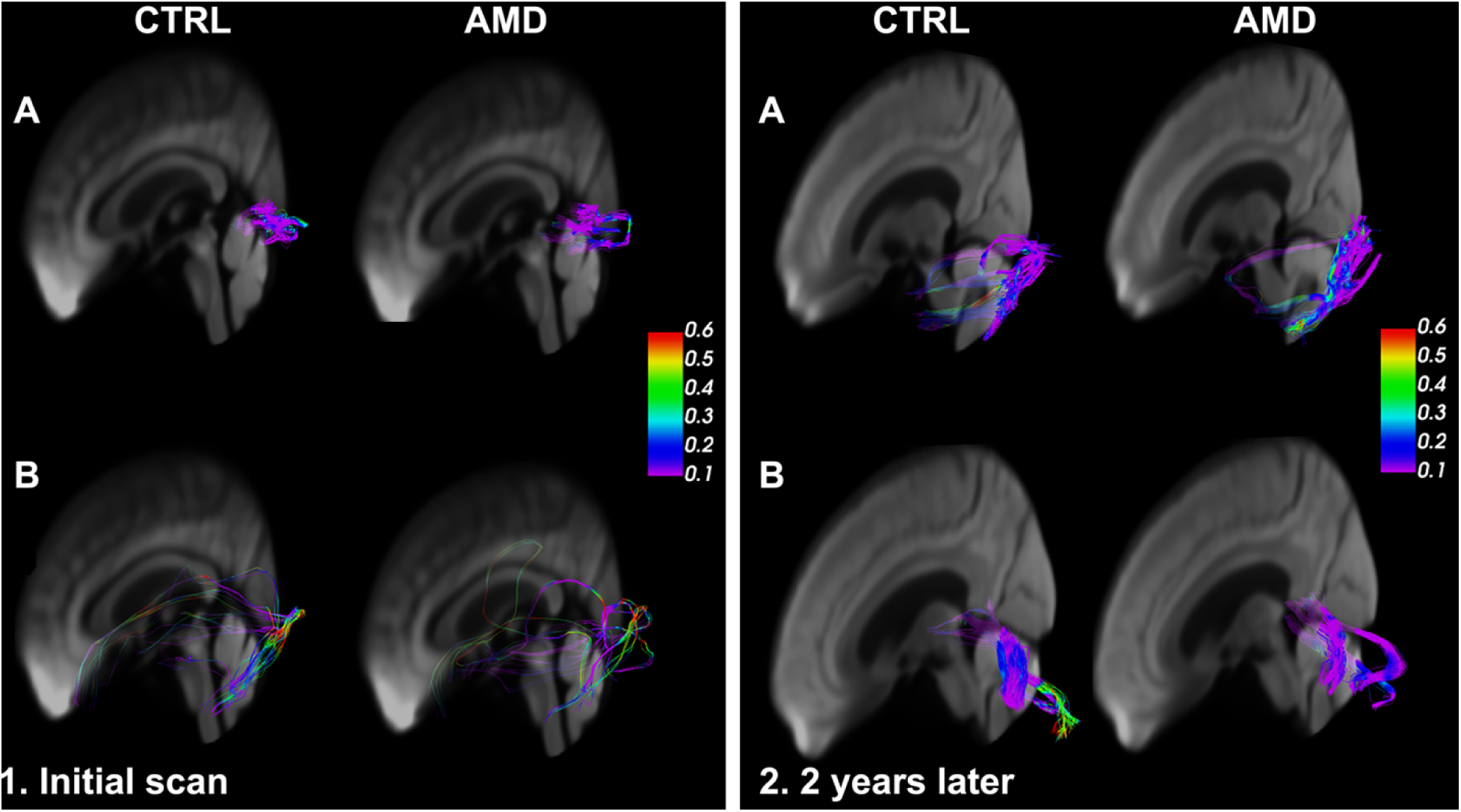
The top connections that differentiate between the groups, colored by FA. 1. Initial visit (A. Lingual Right - Lingual Left; Inferior temporal Right – Superior temporal Left); 2. second visit (A. Left Lingual-Cerebellum Right; B. Lingual Right - Cerebellum Left).

**Figure 4.**
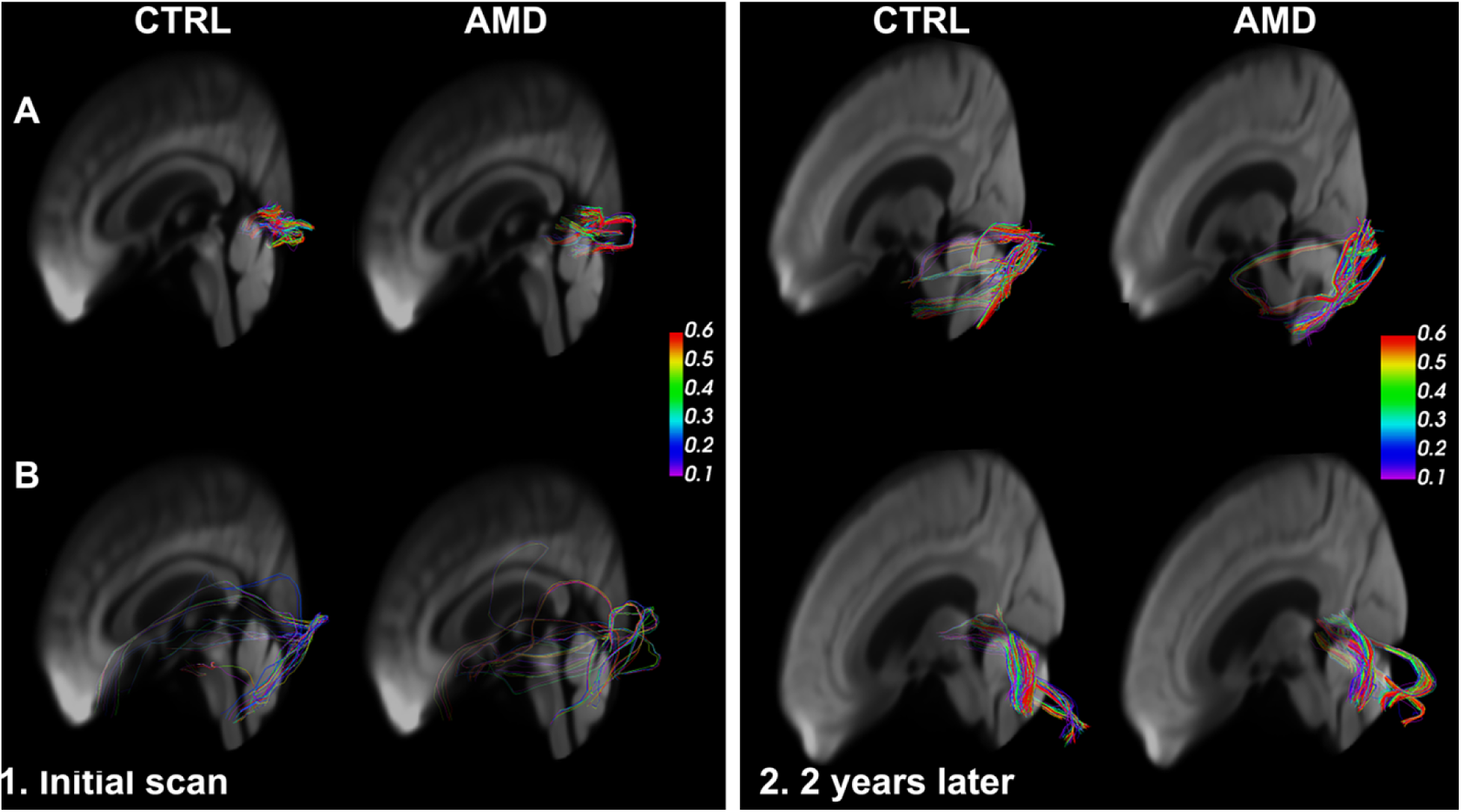
The top connections that differentiate between the groups, colored by the average bundle coherence. 1. At initial visit (A. Lingual Right - Lingual Left; Inferior temporal Right - Superior temporal Left); 2. at the second visit (A. Left Lingual-Cerebellum Right; B. Lingual Right - Cerebellum Left).

**Figure 5.**
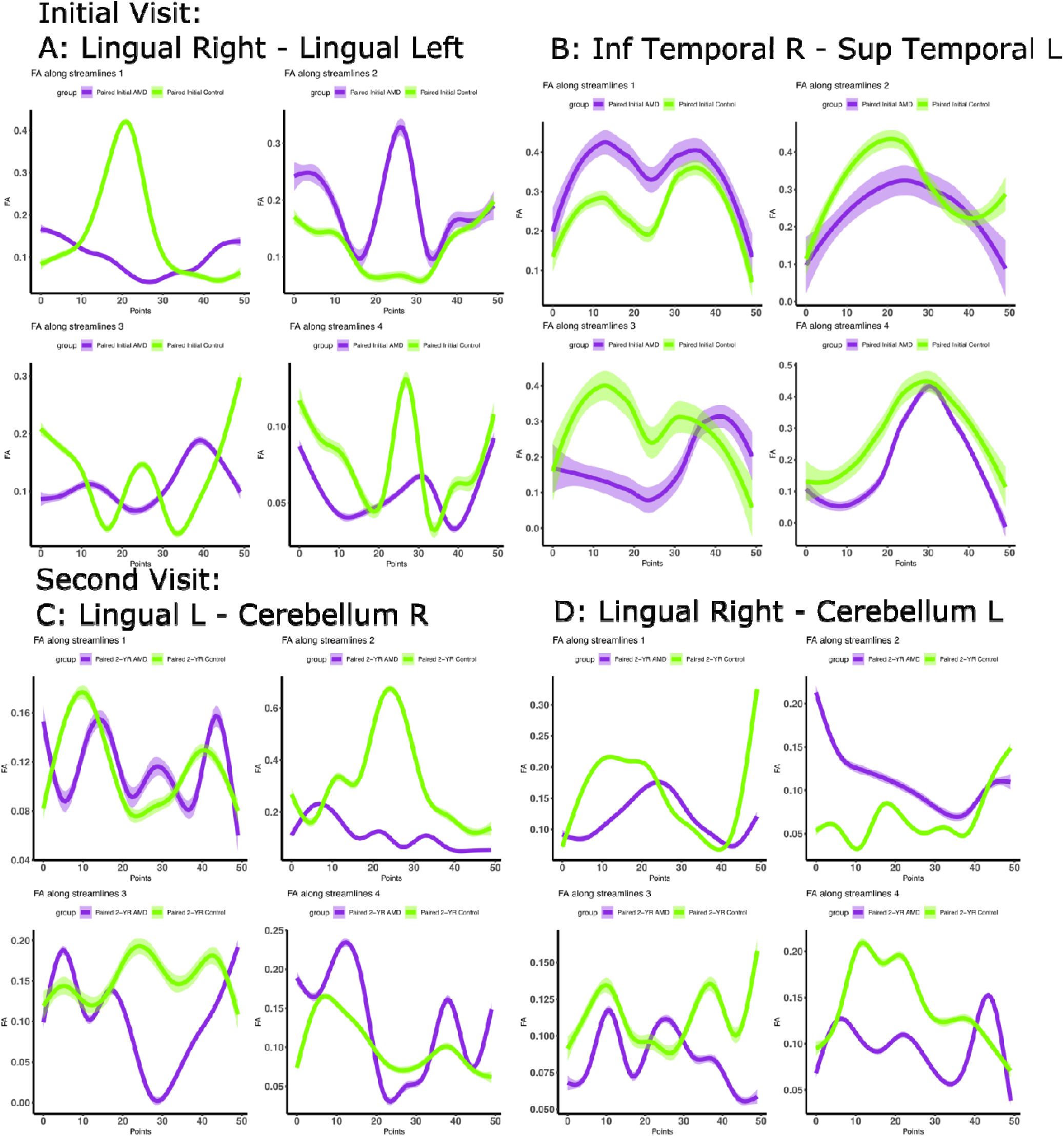
Average FA found along the streamlines of the largest bundles of all tracts corresponding to the top connections determined by TN-PCA. Each bundle was directly compared to its closest equivalent in AMD, as determined by bundle centroid to centroid distance. 1.A. Lingual Right - Lingual Left; Inferior-temporal Right - Superior-temporal Left; 2.A. Left Lingual-Cerebellum Right; B. Lingual Right - Cerebellum Left.

From the tractography results, we were able to observe connectivity changes associated with AMD over time. Overall, there seemed to be a slightly higher number of streamlines in controls than in AMD, and this difference only increased over time. In particular, the pericalcarine cortex had much fewer connections to multiple structures when compared to the Control participants (Table 1). This is particularly significant in the connection from the pericalcarine cortex to the lateral-orbitofrontal cortex, but only at the 2-year acquisition timepoint. Though there are no known associations of changes to the pericalcarine cortex with AMD specifically, there are numerous known changes to the pericalcarine region following vision impairment, including biochemical increases in acetylcholine, glutamate, myo-inositol, increase in gray matter in participants with bilateral anophthalmia [81] and increase in cortical thickness in blind participants [82]. It seems plausible that AMD-related vision loss could cause a progressive atrophy of connections to the pericalcarine gyrus.

Both TN-PCA and subsequent Fisher test indicated a large degree of importance to connections between the lingual gyri of the two hemispheres, and connection between the lingual gyrus and the cerebellum. The FA in controls for the lingual left to right connections was overall much lower than that found in AMD. The FA in controls for the lingual left to right connections was overall much lower than that found in AMD, with this difference only deepening over time. The reasons for this dynamic are as yet unclear as lingual gyrus is linked to processing vision and we would therefore suspect fiber atrophy. However, it should be noted that gliosis and glial scarring can increase FA [83, 84], or that certain regions may try to compensate for the atrophy of others. In participants with retinitis pigmentosa, the lingual gyrus is affected along with the cuneate gyrus and pericalcarine [85]. The lingual gyrus has also been linked to changes to the retinal nerve and Alzheimer’s disease [86, 87]. Resting state fMRI measuring degree centrality, a metric that establishes the number of functional connections at a voxel level as established from functional graphs [15, 88], found that AMD patients have significantly lower degree of centrality in the lingual gyrus compared to controls [89]. Another study showed that a decrease in functional connectivity of the lingual gyrus with the dorsal anterior cortex was associated with Alzheimer’s disease [90]. These studies and our own analysis indicate a connection between retinal health and the lingual gyrus, which could be correlated with declines in memory, whether or not in the concurrent presence of Alzheimer’s disease.

While the cerebellum itself is mostly associated with motor functions, it has for the past few years been thought to be responsible in modulating non-motor language processes and cognitive functions, the right cerebellum lobe in particular [91], though its exact dynamics and involvement remains a subject of hot debate [92]. Still, given AMD’s aforementioned link with linguistic cognitive decline, it is interesting to see significant group differences in the FA and length of the streamlines connecting the cerebellum to lingual gyri, though this effect is mostly seen in the global metric for the connections of the left cerebellum, instead of the right as would be expected due to its more important role in cognitive functioning [93]. Moreover, our study supports an important role for the cerebellar connections, in particular lingual-cerebellar inter-hemispheric connections as they appear to be deteriorating in AMD, possibly in conjunction with visual-spatial learning and memory [94].

Several limitations may impact the interpretation of our results. First, the number of lost participants and overall small sample size limited the power of our comparative analysis. Secondly, our analyses of group differences in brain structure do not incorporate behavioral variables and thus are unbiased; however, this analytical approach means that the regions and connectomes identified here have not been directly linked to specific cognitive or behavioral deficits and our interpretations about their putative role in performance is currently hypothetical. Third, to facilitate recruitment of this older and multimorbid population, the inclusion criteria were fairly broad: any future study should try and set specific criteria as to the type of AMD and the time since initial diagnosis to better compare the development of the condition from a specific timepoint.

Future analysis should focus on the evolving dynamics and number of connections between the superior frontal right and left regions, as it could be a particularly volatile region that sees important changes when AMD is already established. It might be also of great interest to look into the evolution of the connections of the right inferior frontal gyrus and see whether changes in functional connectivity [14] lead to structural reorganization and ultimately, how the brain changes relate to cognitive decline.

Our study shows that visual system changes are associated not only with changes in the brain regions involved in visual processing, but also with brain atrophy in language and memory areas. Some of these changes were only observed at the later time point, which supports the important role of the eye as a potential source for biomarkers in neurodegenerative conditions.

## Conclusion

Our results demonstrate that diffuse volume atrophy and selective DWI-based measures such as FA distinguish AMD from age-matched controls in visual areas, and that overall this effect grows greater over time. Moreover, some regions linked to cognitive performance in memory and language which showed little difference from controls on first analysis showed a more rapid decline when compared again over time. Our longitudinal tensor analysis revealed a clear pattern of AMD-related changes: not only does tractography seeded in visual cortex demonstrate a faster decline in white matter integrity in participants with AMD, but there are more pronounced connectivity changes in AMDs vs. controls in regions that have been linked to language, speech and memory. Identifying specific patterns of regional atrophy and connectopathy may provide greater insight into the mechanisms associated with greater cognitive decline in participants with AMD compared to the general population.

## Acknowledgement

We are grateful to Lucy Upchurch, Chris Petty and Francis Favorini for help setting up, troubleshooting, and maintaining the computational environment.

This work was supported by RF1 AG057895, R01 AG066184, R01AG043438, RF1 AG070149, P30AG028716, P30AG072978, and P30 AG072958. We are grateful to the members of Alzheimer’s Center, Pepper Center, BIAC, the Duke Radiology, and Duke Neurology Departments for generously sharing advice, support and resources needed for the experiments.

## References

1. Klein, R., B.E. Klein, and K.J. Cruickshanks, The prevalence of age-related maculopathy by geographic region and ethnicity. Prog Retin Eye Res, 1999. 18(3): p. 371–89.

2. Wong, T.Y., et al., The natural history and prognosis of neovascular age-related macular degeneration: a systematic review of the literature and meta-analysis. Ophthalmology, 2008. 115(1): p. 116–26.

3. Wong, W.L., et al., Global prevalence of age-related macular degeneration and disease burden projection for 2020 and 2040: a systematic review and meta-analysis. Lancet Glob Health, 2014. 2(2): p. e106–16.

4. Baker, M.L., et al., Early age-related macular degeneration, cognitive function, and dementia: the Cardiovascular Health Study. Arch Ophthalmol, 2009. 127(5): p. 667–73.

5. Nowak, J.Z., Age-related macular degeneration (AMD): pathogenesis and therapy. Pharmacol Rep, 2006. 58(3): p. 353–63.

6. Pennington, K.L. and M.M. DeAngelis, Epidemiology of age-related macular degeneration (AMD): associations with cardiovascular disease phenotypes and lipid factors. Eye Vis (Lond), 2016. 3: p. 34.

7. Wong, T.Y., et al., Is early age-related maculopathy related to cognitive function? The Atherosclerosis Risk in Communities Study. Am J Ophthalmol, 2002. 134(6): p. 828–35.

8. Pham, T.Q., et al., Relation of age-related macular degeneration and cognitive impairment in an older population. Gerontology, 2006. 52(6): p. 353–8.

9. Woo, S.J., et al., Cognitive impairment in age-related macular degeneration and geographic atrophy. Ophthalmology, 2012. 119(10): p. 2094–101.

10. Rong, S.S., et al., Comorbidity of dementia and age-related macular degeneration calls for clinical awareness: a meta-analysis. Br J Ophthalmol, 2019. 103(12): p. 1777–1783.

11. Clemons, T.E., M.W. Rankin, and W.L. McBee, Cognitive impairment in the age-related eye disease study: AREDS report no. 16. Archives of Ophthalmology (Chicago, Ill.: 1960), 2006. 124(4): p. 537–543.

12. Whitson, H.E., et al., Prevalence and patterns of comorbid cognitive impairment in low vision rehabilitation for macular disease. Archives of gerontology and geriatrics, 2010. 50(2): p. 209–212.

13. Zhuang, J., et al., Cerebral white matter connectivity, cognition, and age-related macular degeneration. NeuroImage: Clinical, 2021. 30: p. 102594.

14. Zhuang, J., et al., Language processing in age-related macular degeneration associated with unique functional connectivity signatures in the right hemisphere. Neurobiol Aging, 2018. 63: p. 65–74.

15. Zuo, X.N., et al., Network centrality in the human functional connectome. Cereb Cortex, 2012. 22(8): p. 1862–75.

16. Budd Haeberlein, S., et al., Two randomized phase 3 studies of aducanumab in early Alzheimer’s disease. The Journal of Prevention of Alzheimer’s Disease, 2022. 9(2): p. 197–210.

17. Middleton, L.E. and K. Yaffe, Promising strategies for the prevention of dementia. Archives of neurology, 2009. 66(10): p. 1210–1215.

18. Wittich, W., et al., Effect of Reading Rehabilitation for Age-Related Macular Degeneration on Cognitive Functioning: Protocol for a Nonrandomized Pre-Post Intervention Study. JMIR research protocols, 2021. 10(3): p. e19931.

19. Klaver, C.C., et al., Is age-related maculopathy associated with Alzheimer’s Disease? The Rotterdam Study. Am J Epidemiol, 1999. 150(9): p. 963–8.

20. Keenan, T.D., R. Goldacre, and M.J. Goldacre, Associations between age-related macular degeneration, Alzheimer disease, and dementia: record linkage study of hospital admissions. JAMA Ophthalmol, 2014. 132(1): p. 63–8.

21. Wen, L.Y., et al., Increased risk of Alzheimer’s disease among patients with age-related macular degeneration: A nationwide population-based study. PLoS One, 2021. 16(5): p. e0250440.

22. Curcio, C.A., et al., Aging of the human photoreceptor mosaic: evidence for selective vulnerability of rods in central retina. Invest Ophthalmol Vis Sci, 1993. 34(12): p. 3278–96.

23. Green, W.R. and C. Enger, Age-related macular degeneration histopathologic studies. The 1992 Lorenz E. Zimmerman Lecture. Ophthalmology, 1993. 100(10): p. 1519–35.

24. Sarks, S.H., et al., Early drusen formation in the normal and aging eye and their relation to age related maculopathy: a clinicopathological study. Br J Ophthalmol, 1999. 83(3): p. 358–68.

25. Dentchev, T., et al., Amyloid-beta is found in drusen from some age-related macular degeneration retinas, but not in drusen from normal retinas. Mol Vis, 2003. 9: p. 184–90.

26. Anderson, D.H., et al., A role for local inflammation in the formation of drusen in the aging eye. Am J Ophthalmol, 2002. 134(3): p. 411–31.

27. Johnson, L.V., et al., The Alzheimer’s A beta -peptide is deposited at sites of complement activation in pathologic deposits associated with aging and age-related macular degeneration. Proc Natl Acad Sci U S A, 2002. 99(18): p. 11830–5.

28. Anderson, D.H., et al., Characterization of beta amyloid assemblies in drusen: the deposits associated with aging and age-related macular degeneration. Exp Eye Res, 2004. 78(2): p. 243–56.

29. Biron, K.E., et al., Amyloid triggers extensive cerebral angiogenesis causing blood brain barrier permeability and hypervascularity in Alzheimer’s disease. PLoS One, 2011. 6(8): p. e23789.

30. Hwang, P.H., et al., Longitudinal changes in hearing and visual impairments and risk of dementia in older adults in the United States. JAMA Network Open, 2022. 5(5): p. e2210734–e2210734.

31. Maharani, A., et al., Associations between self-reported sensory impairment and risk of cognitive decline and impairment in the health and retirement study cohort. The Journals of Gerontology: Series B, 2020. 75(6): p. 1230–1242.

32. Zhao, X., et al., Associations of sensory impairment and cognitive function in middle-aged and older Chinese population: The China Health and Retirement Longitudinal Study. Journal of global health, 2021. 11.

33. Yorgason, J.B., et al., The Longitudinal Association of Late-Life Visual and Hearing Difficulty and Cognitive Function: The Role of Social Isolation. Journal of Aging and Health, 2022: p. 08982643211063338.

34. Ge, S., et al., Longitudinal Association Between Hearing Loss, Vision Loss, Dual Sensory Loss, and Cognitive Decline. J Am Geriatr Soc, 2021. 69(3): p. 644–650.

35. Deal, J.A., et al., Relationship of hearing impairment with MRI brain volumes and cognitive decline in the Atherosclerosis Risk in Communities study. Alzheimer’s & Dementia, 2020. 16(e046473).

36. Zuo, X., et al., Relationship between neural functional connectivity and memory performance in age-related macular degeneration. Neurobiology of aging, 2020. 95: p. 176–185.

37. Markl, M. and J. Leupold, Gradient echo imaging. J Magn Reson Imaging, 2012. 35(6): p. 1274–89.

38. Anderson, R.J., et al., Optimizing Diffusion Imaging Protocols for Structural Connectomics in Mouse Models of Neurological Conditions. Front Phys, 2020. 8.

39. Manjón, J.V., et al., Diffusion weighted image denoising using overcomplete local PCA. PloS one, 2013. 8(9): p. e73021.

40. Smith, S.M., Fast robust automated brain extraction. Hum Brain Mapp, 2002. 17(3): p. 143–55.

41. Avants, B.B., et al., Symmetric diffeomorphic image registration with cross-correlation: evaluating automated labeling of elderly and neurodegenerative brain. Med Image Anal, 2008. 12(1): p. 26–41.

42. Yeh, F.C., V.J. Wedeen, and W.Y. Tseng, Generalized q-sampling imaging. IEEE Trans Med Imaging, 2010. 29(9): p. 1626–35.

43. Garyfallidis, E., et al., Dipy, a library for the analysis of diffusion MRI data. Front Neuroinform, 2014. 8: p. 8.

44. Zhang, S. and K. Arfanakis, Evaluation of standardized and study-specific diffusion tensor imaging templates of the adult human brain: Template characteristics, spatial normalization accuracy, and detection of small inter-group FA differences. Neuroimage, 2018. 172: p. 40–50.

45. Anderson, R.J., et al., Small Animal Multivariate Brain Analysis (SAMBA) - a High Throughput Pipeline with a Validation Framework. Neuroinformatics, 2019. 17(3): p. 451–472.

46. Penny, W.D., et al., Statistical parametric mapping: the analysis of functional brain images. 2011: Elsevier.

47. Nichols, K., False Discovery Rate procedures, in Statistical Parametric Mapping, K. Friston, et al., Editors. 2007, Academic Press: London. p. 246–252.

48. Tuch, D.S., Q-ball imaging. Magn Reson Med, 2004. 52(6): p. 1358–72.

49. Qi, X., S. Zhang, and K. Arfanakis, IIT Human Brain Atlas: enhancement of T1-weighted Template In: Tissue Probability Maps and Gray Matter Atlas. Proc. Int. Soc. for Magn. Reson. In Med.(ISRMRM), 2017. 4688.

50. Zhang, Z., et al., Tensor network factorizations: Relationships between brain structural connectomes and traits. Neuroimage, 2019. 197: p. 330–343.

51. Badea, A., et al., Identifying Vulnerable Brain Networks in Mouse Models of Genetic Risk Factors for Late Onset Alzheimer’s Disease. Front Neuroinform, 2019. 13: p. 72.

52. Baran, T.M., et al., Brain structural connectomes indicate shared neural circuitry involved in subjective experience of cognitive and physical fatigue in older adults. Brain imaging and behavior, 2020. 14(6): p. 2488–2499.

53. Wang, L., Z. Zhang, and D. Dunson, Symmetric bilinear regression for signal subgraph estimation. IEEE Transactions on Signal Processing, 2019. 67(7): p. 1929–1940.

54. Garyfallidis, E., et al., Recognition of white matter bundles using local and global streamline-based registration and clustering. Neuroimage, 2018. 170: p. 283–295.

55. Garyfallidis, E., et al., QuickBundles, a Method for Tractography Simplification. Front Neurosci, 2012. 6: p. 175.

56. Meesters, S.S., et al. Cleaning output of tractography via fiber to bundle coherence, a new open source implementation. 2016.

57. Chandio, B.Q., et al., Bundle analytics, a computational framework for investigating the shapes and profiles of brain pathways across populations. Sci Rep, 2020. 10(1): p. 17149.

58. Cohen, J., F Tests of Variance Proportions in Multiple Regression/Correlation Analysis, in Statistical Power Analysis for the Behavioral Sciences, J. Cohen, Editor. 1977, Academic Press. p. 407–453.

59. Coleman, H.R., et al., Age-related macular degeneration. Lancet, 2008. 372(9652): p. 1835–45.

60. Palejwala, A.H., et al., Anatomy and white matter connections of the lingual gyrus and cuneus. World Neurosurgery, 2021. 151: p. e426–e437.

61. Fusar-Poli, P., et al., Functional atlas of emotional faces processing: a voxel-based meta-analysis of 105 functional magnetic resonance imaging studies. Journal of Psychiatry and Neuroscience, 2009. 34(6): p. 418–432.

62. Grill-Spector, K. and R. Malach, The human visual cortex. Annual review of neuroscience, 2004. 27(1): p. 649–677.

63. Nomi, J.S., et al., On the neural networks of empathy: A principal component analysis of an fMRI study. Behavioral and Brain Functions, 2008. 4(1): p. 1–13.

64. Lepore, N., et al., Brain structure changes visualized in early- and late-onset blind subjects. Neuroimage, 2010. 49(1): p. 134–40.

65. Burton, H., Visual cortex activity in early and late blind people. J Neurosci, 2003. 23(10): p. 4005–11.

66. Amedi, A., et al., The Occipital Cortex in the Blind:Lessons About Plasticity and Vision. Current Directions in Psychological Science, 2005. 14(6): p. 306–311.

67. Barris, R.W. and H.R. Schuman, [Bilateral anterior cingulate gyrus lesions; syndrome of the anterior cingulate gyri]. Neurology, 1953. 3(1): p. 44–52.

68. Leech, R. and D.J. Sharp, The role of the posterior cingulate cortex in cognition and disease. Brain, 2014. 137(Pt 1): p. 12–32.

69. du Boisgueheneuc, F., et al., Functions of the left superior frontal gyrus in humans: a lesion study. Brain, 2006. 129(Pt 12): p. 3315–28.

70. Binder, J.R., Current Controversies on Wernicke’s Area and its Role in Language. Curr Neurol Neurosci Rep, 2017. 17(8): p. 58.

71. Blank, S.C., et al., Speech production: Wernicke, Broca and beyond. Brain, 2002. 125(Pt 8): p. 1829–38.

72. Ohyama, M., et al., Role of the nondominant hemisphere and undamaged area during word repetition in poststroke aphasics. A PET activation study. Stroke, 1996. 27(5): p. 897–903.

73. Counts, S.E., et al., Differential expression of synaptic proteins in the frontal and temporal cortex of elderly subjects with mild cognitive impairment. J Neuropathol Exp Neurol, 2006. 65(6): p. 592–601.

74. Gross, C.G., How inferior temporal cortex became a visual area. Cereb Cortex, 1994. 4(5): p. 455–69.

75. Karnath, H.O., New insights into the functions of the superior temporal cortex. Nat Rev Neurosci, 2001. 2(8): p. 568–76.

76. Scheff, S.W., et al., Synaptic loss in the inferior temporal gyrus in mild cognitive impairment and Alzheimer’s disease. J Alzheimers Dis, 2011. 24(3): p. 547–57.

77. Baseler, H.A., et al., Large-scale remapping of visual cortex is absent in adult humans with macular degeneration. Nat Neurosci, 2011. 14(5): p. 649–55.

78. He, B.J., et al., Breakdown of functional connectivity in frontoparietal networks underlies behavioral deficits in spatial neglect. Neuron, 2007. 53(6): p. 905–18.

79. Capotosto, P., et al., Frontoparietal cortex controls spatial attention through modulation of anticipatory alpha rhythms. J Neurosci, 2009. 29(18): p. 5863–72.

80. Wurm, M.F. and A. Caramazza, Distinct roles of temporal and frontoparietal cortex in representing actions across vision and language. Nat Commun, 2019. 10(1): p. 289.

81. Coullon, G.S., et al., Neurochemical changes in the pericalcarine cortex in congenital blindness attributable to bilateral anophthalmia. J Neurophysiol, 2015. 114(3): p. 1725–33.

82. Georgy, L., et al., Changes in peri-calcarine cortical thickness in blindsight. Neuropsychologia, 2020. 143: p. 107463.

83. Budde, M.D., et al., The contribution of gliosis to diffusion tensor anisotropy and tractography following traumatic brain injury: validation in the rat using Fourier analysis of stained tissue sections. Brain, 2011. 134(8): p. 2248–2260.

84. Harris, N.G., et al., Bi-directional changes in fractional anisotropy after experiment TBI: disorganization and reorganization? Neuroimage, 2016. 133: p. 129–143.

85. Rita Machado, A., et al., Structure-function correlations in Retinitis Pigmentosa patients with partially preserved vision: a voxel-based morphometry study. Scientific reports, 2017. 7(1): p. 11411–11411.

86. Shen, Y., et al., The attenuation of retinal nerve fiber layer thickness and cognitive deterioration. Front Cell Neurosci, 2013. 7: p. 142.

87. Shi, Z., et al., Greater attenuation of retinal nerve fiber layer thickness in Alzheimer’s disease patients. J Alzheimers Dis, 2014. 40(2): p. 277–83.

88. Wang, J., X. Zuo, and Y. He, Graph-based network analysis of resting-state functional MRI. Front Syst Neurosci, 2010. 4: p. 16.

89. Huang, R., et al., Altered Spontaneous Brain Activity Patterns in Patients with Age-related Macular Degeneration: A Resting-State fMRI Study Running head: Functional Connectivity Density alterations in age-related macular degeneration patients. 2021, Research Square.

90. Liu, X., et al., Decreased functional connectivity between the dorsal anterior cingulate cortex and lingual gyrus in Alzheimer’s disease patients with depression. Behav Brain Res, 2017. 326: p. 132–138.

91. Silveri, M.C. and S. Misciagna, Language, memory, and the cerebellum. Journal of neurolinguistics, 2000. 13(2-3): p. 129–143.

92. Mariën, P., et al., Consensus paper: language and the cerebellum: an ongoing enigma. The Cerebellum, 2014. 13(3): p. 386–410.

93. Murdoch, B.E., The cerebellum and language: historical perspective and review. Cortex, 2010. 46(7): p. 858–868.

94. Doyon, J., et al., Experience-dependent changes in cerebellar contributions to motor sequence learning. Proceedings of the National Academy of Sciences, 2002. 99(2): p. 1017–1022.

