## Supplementary figures and images for "Age-related Macular Degeneration is associated with faster rates of structural brain changes and widespread differences in connectivity"

### Supplementary Figure

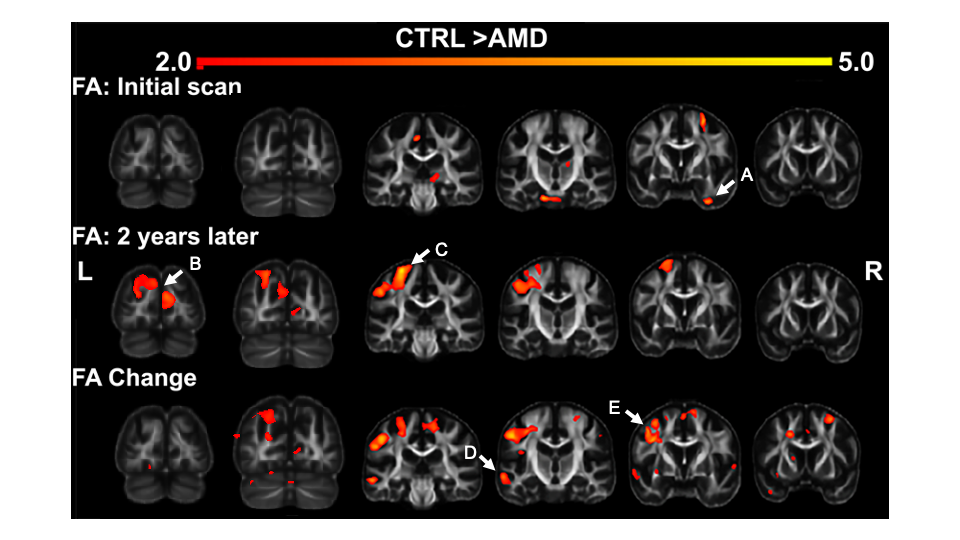
